# Activation of G-protein coupled receptors is thermodynamically linked to lipid solvation

**DOI:** 10.1101/2020.08.20.259077

**Authors:** Alison N. Leonard, Edward Lyman

## Abstract

Preferential lipid solvation of the G-protein coupled A_2A_ adenosine receptor (A_2A_R) is evaluated from 35 *μ*sec of all-atom molecular dynamics simulation. A coarse-grained transition matrix algorithm is developed to overcome slow equilibration of the first solvation shell, obtaining statistically robust estimates of the free energy of solvation by different lipids for the receptor in different activation states. Results indicate preference for solvation by unsaturated chains, which favors the active receptor. A model for lipid-dependent GPCR activity is proposed in which the chemical potential of lipids in the bulk membrane modulates receptor activity. The enthalpy and entropy associated with moving saturated vs. unsaturated lipids from bulk to A_2A_R’s first solvation shell are compared. In the simulated mixture, saturated chains are disordered (i.e., obtain a favorable entropic contribution) when partitioning to the receptor surface, but this is outweighed by a favorable enthalpic contribution for unsaturated chains to occupy the first solvation shell.

## Introduction

Lipid composition is highly differentiated among different cell types (Řezanka et al., 2018; Pradas et al., 2018), between the membranes within a single cell (Doralicia et al., 2019), and even between the leaflets of a single membrane (Lorent et al., 2020). These distinct lipid environments are known to influence the function of membrane proteins such as G-protein coupled receptors (GPCRs) (Periole, 2017) — transmembrane proteins that comprise the largest class of proteins in the human genome and are important drug targets (Katritch et al., 2013). Understanding lipid-dependent activity of GPCRs might therefore yield new therapeutic strategies based on tissue- or context-specific pharmacological targeting.

Tight, site-specific interactions between lipids and GPCRs have been detected using x-ray diffraction (Yeagle, 2014), electron microscopy (Gonen et al., 2005), and high resolution mass spectrometry (Bolla et al., 2019; Montenegro et al., 2017), and so mechanisms have tended to focus on specific cholesterol (Cang et al., 2013; Hanson et al., 2008; Lee et al., 2013; McGraw et al., 2019; Rouviere et al., 2017; Gutierrez et al., 2016; O’Malley et al., 2011; Lee & Lyman, 2013) and lipid (Javanainen et al., 2019; Grossfield et al., 2006; Neale et al., 2015; Song et al., 2019; Naranjo et al., 2016) interactions. In contrast, resolving lipid solvation of a transmembrane protein (i.e., weaker but still preferential interactions) presents a challenge for experiment, requiring labeled protein (Loura et al., 2010; Soubias & Gawrisch, 2005) or lipid probes (Marsh et al., 1982; Marsh, 1990). The best studied member of the family is rhodopsin, for which NMR (Soubias & Gawrisch, 2005; Soubias & Gawrisch, 2012; Soubias et al., 2008; Feller et al., 2003), simulation (Grossfield et al., 2006), and theoretical (Botelho et al., 2002; Huber et al., 2004) work suggest that the negative intrinsic curvature of polyunsaturated fatty acid (PUFA)-containing lipids shifts the receptor toward active conformations. Javanainen, et al. found in simulations of A_2A_R that PUFA chains in the first solvation shell mediate partitioning of the receptor into ordered lipid phases, following experimental partitioning measurements in phase separated GUVs reported by Guttierrez and Malmstadt (Javanainen et al., 2019; Gutierrez et al., 2019). Yang and Lyman also reported preferential solvation of A_2A_R by the monounsaturated chains of dioleyoyl phospholipids relative to saturated lipids (Yang & Lyman, 2019).

But is the preference for acyl chain unsaturation coupled with receptor activity? In the simplest model for GPCR activation, the receptor is in a conformational equilibrium between an inactive “R” state and an active “R*” state. In this model, anything which shifts the R/R* equilibrium will modulate activity (Leff, 1995). Thus if the R and R* states recruit different lipid environments, changes in lipid solvation will influence the R/R* equilibrium and modulate activity. In more complex models of receptor function, the receptor can adopt multiple states which couple to different pathways (e.g., different G-proteins or arrestins) (Nygaard et al., 2013), and the extent to which the lipid environment shifts the balance of these multiple states will modulate signaling output.

The present work addresses this question by all-atom simulations of the inactive and partially-active A_2A_R in a mixture of dipalmitoylphosphatidylcholine (DPPC), cholesterol, and dioleoylphosphatidylcholine (DOPC). Statistically significant differences in lipid solvation between the inactive and partially-active (agonist bound, but without G-protein) receptors are found. The effect of these differences is to shift the R↔R* equilibrium, with the active receptor recruiting on average three more unsaturated lipids to its first solvation shell than the inactive receptor. Thermodynamic analysis of the first solvation shell indicates this represents a difference in free energy of about −0.32 kcal/mol. The consequences of state-dependent lipid solvation are analyzed within the context of a thermodynamic model that incorporates ligand binding, the R↔R* conformational equilibrium, and lipid solvation.

Fluctuations in the lipid composition of the first solvation shell are significant even during 15-*μ*sec all-atom simulations (Yang & Lyman, 2019), but statistically reliable estimates of the lipid solvation free energy require determining the equilibrium composition of the first shell with high precision. To address this problem, a new coarse-grained transition matrix (CGTM) algorithm is developed specifically to equilibrate the lipid environment around integral membrane proteins. The CGTM method permits a rigorous assessment of statistical uncertainty, and can readily be applied to simulations of other membrane proteins.

## Results

### Equilibrium distribution of the first lipid solvation shell of A_2A_R by the CGTM approach

The 15 *μ*sec inactive and partially-active trajectories in the present work were reported previously (Yang & Lyman, 2019), It was found that the first solvation shell of A_2A_R is enriched in unsaturated DOPC relative to bulk concentration (see Figure 1A). The previous report included a histogram of first-shell lifetimes, which ranged from 5 nsec to the length of the 15-*μ*sec simulation in the case of three cholesterols. The average first-shell lifetime for all three lipid types was reported to be ∼100 nsec.

**Figure 1.**
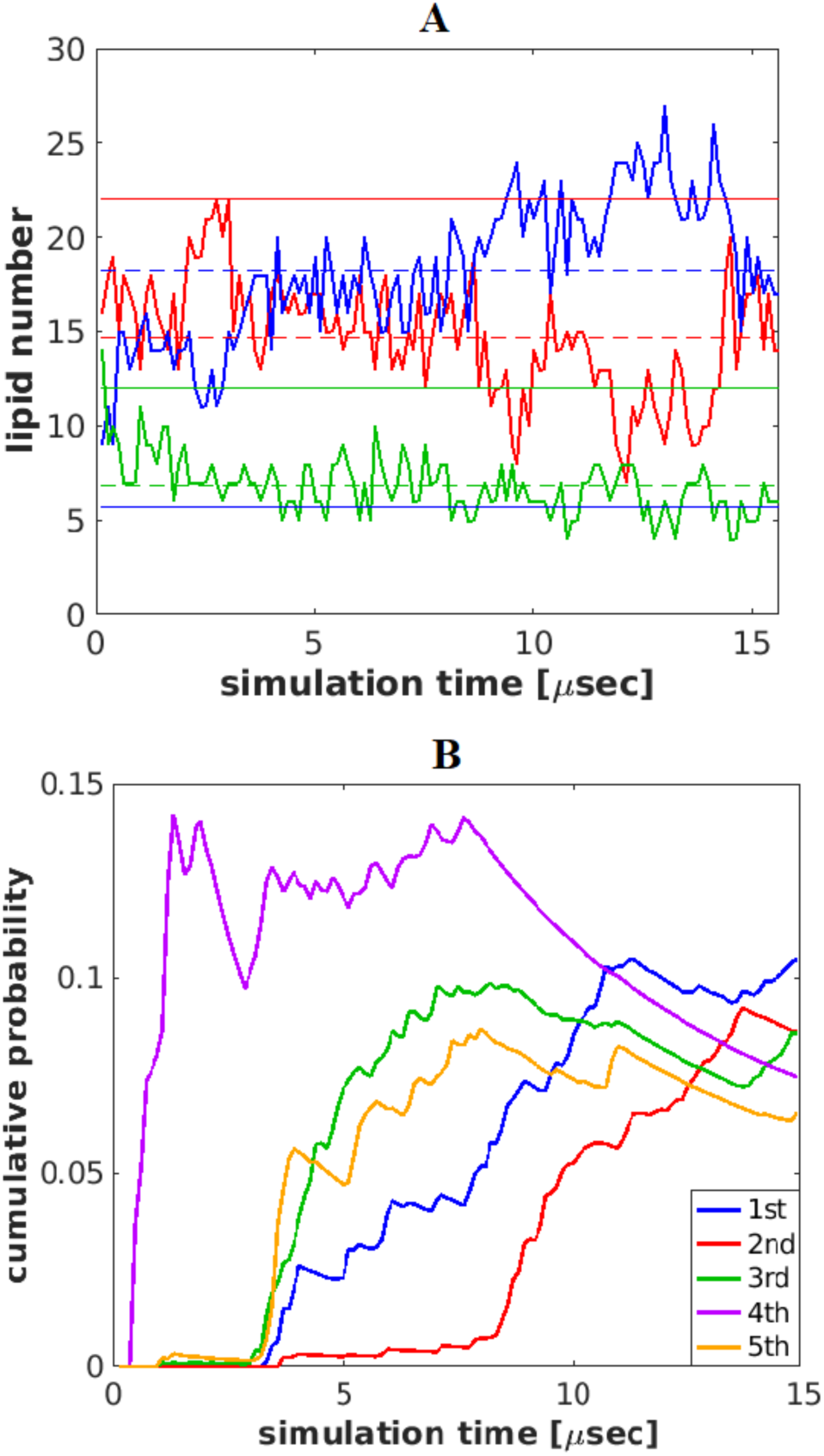
First lipid solvation shell of receptor does not converge over a 15 *μ*sec all-atom simulation. Panel A: Composition of the first lipid solvation shell; agonist bound receptor (similar results are obtained for the inactive receptor). The number of each lipid type in the first solvation shell is obtained by the Voronoi analysis described in Methods. Red, DPPC; blue, DOPC; green, cholesterol. Horizontal dashed lines show the average over entire simulation of the first shell, solid lines show the bulk composition. Both leaflets are included. Panel B: Probabilities of the 5 most populated states over time; agonist-bound receptor. Probability is integrated over time with 125 evenly-spaced sampling events. The following states are plotted with DPPC/DOPC/Chol compositions given as percentages: 1^st^ 35/50/15; 2^nd^ 30/55/15; 3^rd^ 40/45/15; 4^th^ 40/40/20; 5^th^ 35/45/20.

The scale of fluctuations in Figure 1A suggests significant uncertainty in the average first shell lipid composition. This motivated the construction of a CGTM to compute the equilibrium first shell occupancy. Coarse-grained states were defined by binning the first shell occupancies in increments of 5 mol % of each lipid type (roughly equal to two lipids); the partially-active receptor occupied 63 and the inactive receptor 58 distinct states in the respective 15-*μ*sec simulations. The need for improved sampling is apparent in Figure 1B, which plots the cumulative probabilities of the five most populated states in the partially-active trajectory, together accounting for nearly 50% of the total probability. Equilibration of these states is not achieved by the end of the 15 *μ*sec simulation — note the nonzero slopes at the end of the simulation, indicating that the occupancy of these states is still changing in time. The fact that one-half of the total probability is accounted for by only five states also raises the question of whether the sampling is broad enough in the 15 *μ*sec trajectories.

To address sampling limitations, the CGTM approach was implemented as described in Methods. A total of 144 trajectories per receptor state, each 50 nsec in duration (72 taken from non-overlapping segments of the 15-*μ*sec trajectory and 72 seeded from under-sampled states) were used to estimate the transition matrix. The state populations predicted by this transition matrix (denoted *π*^Eq^) are reported in Figure 2A (top panel) for the partially active receptor. For comparison, the average occupancies of the same states were obtained directly from the 15-*μ*sec trajectory (denoted *π*^15*μ*s^) and are shown in Figure 2A (bottom panel). The same comparison is made for the inactive receptor in Fig. 2B. Both states preferentially recruit unsaturated DOPC over saturated DPPC despite the fourfold elevation of DPPC in bulk, as previously reported (Yang & Lyman, 2019). See Materials and Methods for a complete description of molecular dynamics simulations.

**Figure 2.**
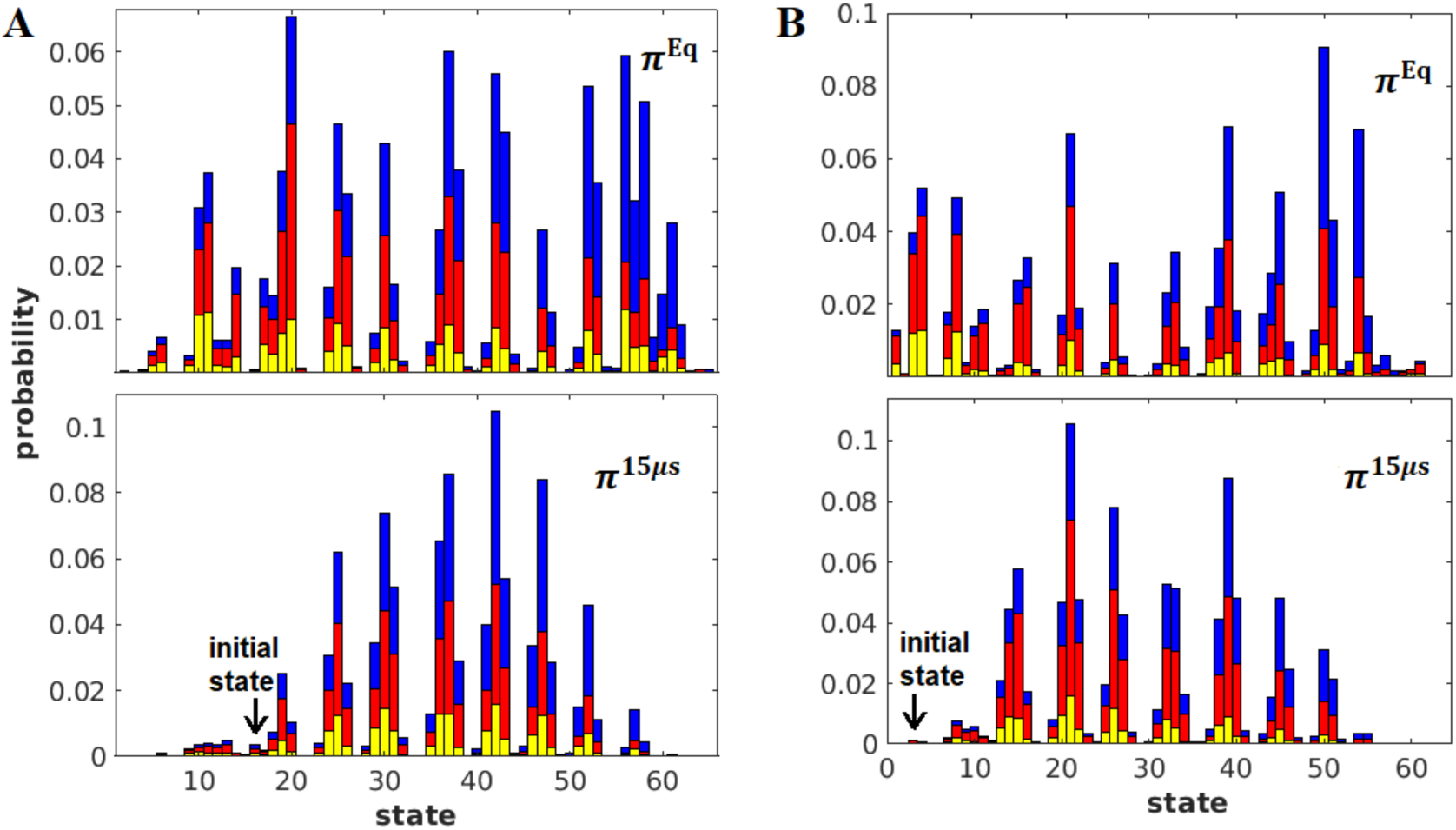
Comparison of occupation probabilities *π*^Eq^ computed from CGTM, and raw occupancies *π*^15*μ*s^ computed from 15-*μ*sec trajectory. Blue: DOPC; red: DPPC; yellow: cholesterol. Each state is a unique ratio of DPPC/DOPC/Cholesterol, binned in 5 mol % increments. Panels A (top and bottom), partially active receptor. Panels B (top and bottom), inactive receptor.

Comparison of the top and bottom panels of Figures 2A and 2B shows that the distributions obtained from the CGTM are broader, indicating improved sampling of states that were under-sampled in the 15-*μ*sec trajectories. However, the average composition (that is, the average ratio of DPPC/DOPC/cholesterol) of the first shell obtained from *π*^Eq^ and *π*^15*μ*s^ (Table 1) are strikingly similar. The fact that this is the case for simulations of both receptor states suggests that this is not a coincidence, and indicates that if the “brute force” simulations were run longer, A_2A_R would sample a broader distribution of solvation states, but that the overall composition of the first shell would not change.

**Table 1.** Fraction of each lipid type in first solvation shell vs. bulk, and free energies to transfer from bulk to first shell. Raw first-shell occupancies from 15-*μ*sec simulations (*π*^15μs^) and equilibrium distribution computed by the CGTM approach (*π*^Eq^), compared with bulk distribution (*π*^Bulk^). Free energies obtained from *π*^Eq^ values (Eqs. 1 and 2). Errors in *π*^Eq^ utilize convergence of probabilities to within 1% (See Materials and Methods, Supplemental Figure 6-1).

|  |  | DPPC | DOPC | Chol |  |
| --- | --- | --- | --- | --- | --- |
| Agonist | $\pi^{15\mu s}$ | 36.9% | 45.9% | 17.2% | |
| | $\pi^{\text{Eq}}$ | 35.9±0.4% | 46.7±0.5% | 17.4±0.2% | |
| | $\pi^{\text{Bulk}}$ | 55.5% | 14.3% | 30.2% | $\Delta\Delta G$ (Eqs. 1 & 2)<br>DOPC – DPPC |
| | $\Delta G$, Eq. 1<br>[kcal/mol] | 0.24±0.01 | -0.68±0.01 | 0.33±0.01 | -0.92±0.01 |
| Antagonist | $\pi^{15\mu s}$ | 49.8% | 37.3% | 12.9% | |
| | $\pi^{\text{Eq}}$ | 46.7±0.5% | 38.8±0.4% | 14.5±0.1% | |
| | $\pi^{\text{Bulk}}$ | 55.3% | 14.8% | 30.0% | $\Delta\Delta G$ (Eqs. 1 & 2)<br>DOPC – DPPC |
| | $\Delta G$, Eq. 1<br>[kcal/mol] | 0.06±0.01 | -0.54±0.01 | 0.49±0.01 | -0.60±0.01 |

The free energy difference to bring a lipid from bulk into the first shell can also be estimated directly from *π*^Eq^ of each lipid type in the first shell vs. bulk:

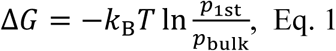

where *p*_1st_ and *p*_bulk_ are the fractional occupancies of each lipid in the 1^st^ shell or in bulk, and Δ*G* is computed for DOPC and DPPC separately. The preference of the receptor for unsaturated lipids can then be quantified by the difference in Δ*G* for the two types of phospholipids:

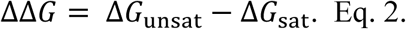

A ne gative ΔΔ*G* represents preference for unsaturated DOPC and a positive ΔΔ*G* preference for saturated DPPC.

ΔΔ*G* for both receptor states are given in Table 1. The partially-active receptor recruits a first solvation shell that is more enriched in DOPC than the inactive receptor. Both receptor states recruit a solvation shell that is depleted in cholesterol relative to bulk. These observations can be made with high confidence thanks to the convergence of first shell occupancies afforded by the CGTM (See Materials and Methods, Supplemental Figures 6-1 and 6-2). This suggests a link between lipid solvation and protein function, which is revisited in the Discussion.

Well-converged lipid solvation was obtained for this system with only 2.4 *μ*sec of total simulation time: (number of initial states) × (number of trajectories per state) × (duration of each trajectory) = 16 × 5 × 30 ns = 2.4 *μ*sec (See Supplemental Figures 6-1 and 6-2). This is less than 1/5 the duration of the 15 *μ*sec simulations which never reached clear convergence. Moreover, the CGTM approach is naively parallel — all data used to obtain the results presented here were obtained in a single day on a commodity supercomputing resource.

The CGTM model can be visualized as the exchange of probability between different solvation states, drawing the transition matrix as a network of states, with the transition probabilities as the links between states as shown in Figure 3. The network reveals a limited number of “bridge” states that connect states with elevated DPPC to states with elevated DOPC. Figure 2 show that both 15 *μ*sec simulations were initialized in relatively underpopulated states. The limited number of “bridge” states suggests why equilibration of the first solvation shell requires such a long brute force simulation time — the system can cycle among elevated DPPC states, for example, for some time before crossing the bridge to sample elevated DOPC states. The network representation also shows how a similar first shell composition is obtained by the long single trajectories and the CGTM method, despite distinct underlying state distributions. Once the long trajectory crosses into the elevated DOPC states (lower left of the panels in Figure 3), it is likely to remain there for an extended period of time, suggesting that the correspondence between the long trajectories and the CGTM model is specific to the topology of the transition matrix, which in turn is likely specific to the membrane composition and perhaps also the protein. It seems impossible to predict this outcome in advance, underlining the value of the CGTM approach.

**Figure 3.**
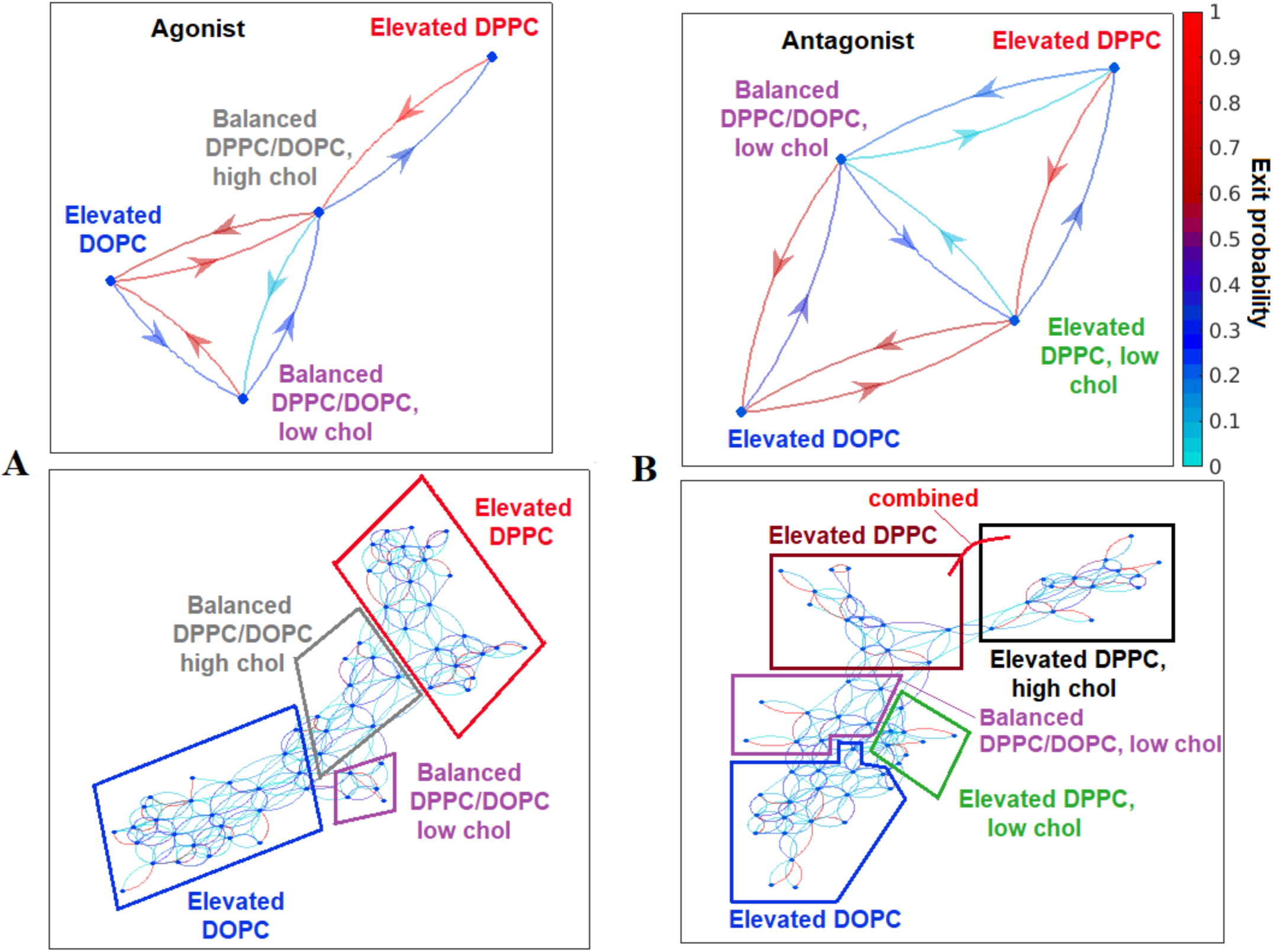
CGTM as a network shows flow among first solvation shell configurations. Top Panels: State space is mapped onto four coarse-grained states based on shape of the map for all states and state characteristics. Bottom Panels: Map of transition flow for the entire state space. Transition probabilities to exit a state are shown; the probability of returning to the same state is omitted. Panels A (top and bottom), partially active receptor. Panels B (top and bottom), inactive receptor.

### Thermodynamic modeling of acyl chain solvation

An independent assessment of lipid solvation was determined by estimating the (Gibbs) free energy associated with bringing a single lipid of either type (DOPC or DPPC) from bulk to the receptor’s first solvation shell, based on separate calculations of the enthalpic (Δ*H*) and entropic (Δ*S*) contributions:

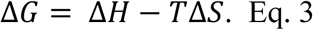

In estimating the Δ*H* term in Eq. 3, only dihedral and non-bonded energies of the acyl chains are considered, since the headgroups of DOPC and DPPC are identical. Details of the calculation are given in Materials and Methods.

Differences in the conformational entropy of the hydrocarbon chains are expected to be an important component of the lipid solvation free energy. The chain conformational entropy Δ*S* is estimated from the Gibbs entropy:

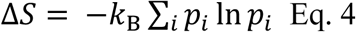

where *k*_B_ is the Boltzmann constant and *p*_*i*_ is the observed probability of configuration *i* of the chain. Following Grossfield, et al. (Grossfield et al., 2006), the configurations are enumerated using a rotational isomeric states model in which each dihedral is assigned to a discrete state based on symmetry. For example, a C-C-C-C dihedral could be (t)rans, (g^+^)auche, or (g^-^)auche. A configuration is defined by the sequence of dihedral states, 13 for a palmitoyl chain and 15 for an oleoyl chain, differing from the previous work which grouped sets of three neighboring dihedrals into a single state to reduce the size of the configuration space. Despite the large state space, good convergence is seen for the resulting estimates of *S* (See Materials and Methods, Supplemental Figure 4-3).

**Figure 4.**
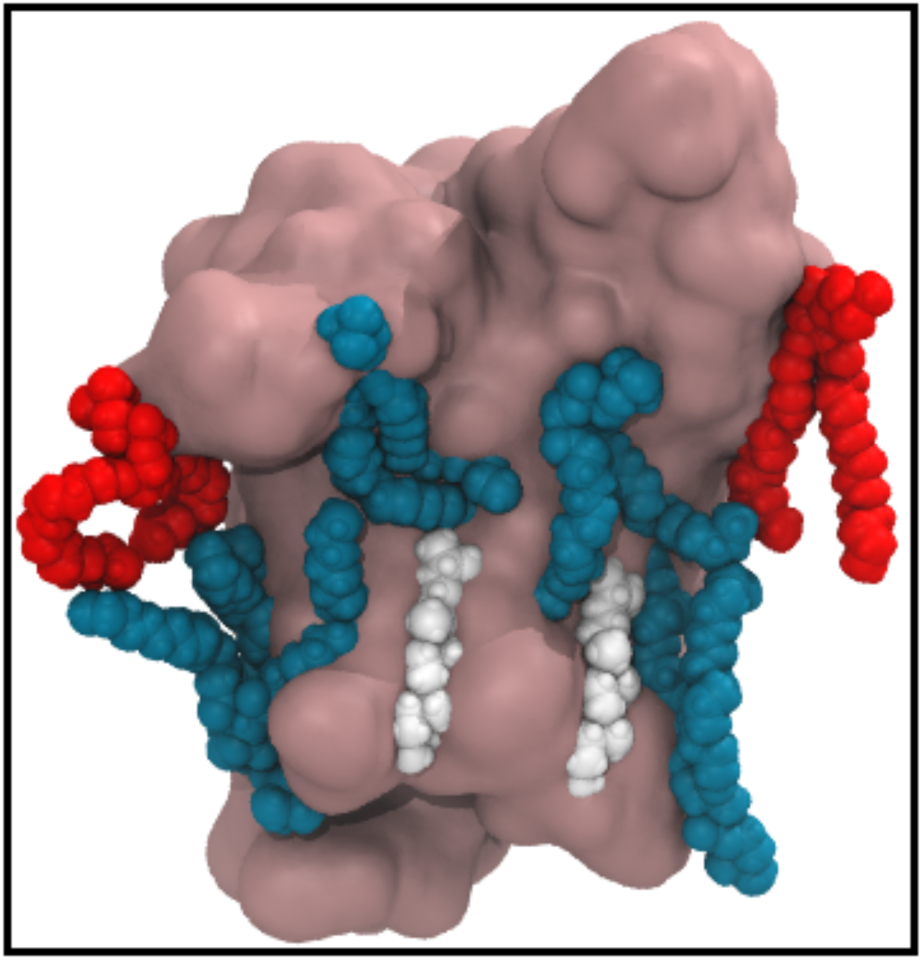
Lipid tails conforming to the receptor surface. Red, DPPC; blue, DOPC; white, cholesterol. Note the conformation of the acyl chains, many of which are contorted as they conform to the receptor surface. Only a subset of first shell lipids are shown for clarity. Partially-active receptor.

Table 2 gives Δ*H* and − *T*Δ*S* for each lipid type. They are defined so that negative values indicate a favorable thermodynamic contribution for that lipid to transfer from the bulk to the receptor surface. The entropic contribution is favorable for both lipids, but more so for DPPC. In the simulated mixture, the bulk lipids are more ordered than those at the protein surface, and this is more pronounced for DPPC. However, all 3 enthalpic contributions favor solvation by DOPC (DOPC Δ*H* are either more negative or are positive but smaller in magnitude than for DPPC). Taken together, the thermodynamics of the hydrocarbon chains yield a favorable free energy for DOPC to partition to the receptor surface in both activation states, but more so for the partially-active state. The thermodynamic analysis is consistent with the partitioning free energies that are obtained directly from relative probabilities for each lipid to be in bulk vs. at the receptor surface, as reported in Table 1. In that case, solvation by DOPC is again favorable, and more so for the partially-active receptor. Absolute numerical agreement is not expected, given that thermodynamic analysis only includes contributions from the hydrocarbon chains. Details of the calculations of Δ*H* and *S* are given in Materials and Methods.

**Table 2.** Contributions to ΔΔ*G* to bring a lipid molecule from bulk to first solvation shell.

| | Bulk → First Shell<br>Contribution [kcal/mol] | DOPC | DPPC | |
| --- | --- | --- | --- | --- |
| Agonist | $\Delta H$, dihedrals | $0.055 \pm 0.02$ | $0.187 \pm 0.02$ | |
| | $\Delta H$, Lennard-Jones | $14.85 \pm 0.06$ | $15.24 \pm 0.07$ | |
| | $\Delta H$, Coulomb | $-6.738 \pm 0.05$ | $-4.851 \pm 0.06$ | |
| | $-T\Delta S$, dihedrals | <u>$+ -0.543 \pm 0.005$</u> | <u>$+ -2.525 \pm 0.007$</u> | $\Delta\Delta G$ (Eqs. 3 & 2)<br>DOPC - DPPC |
| | $\Delta G$, sum, Eq. 3 | $7.62 \pm 0.08$ | $8.05 \pm 0.09$ | $-0.43 \pm 0.11$ |
| Antagonist | $\Delta H$, dihedrals | $0.026 \pm 0.02$ | $0.115 \pm 0.01$ | |
| | $\Delta H$, Lennard-Jones | $13.80 \pm 0.04$ | $14.97 \pm 0.05$ | |
| | $\Delta H$, Coulomb | $-5.682 \pm 0.07$ | $-4.827 \pm 0.06$ | |
| | $-T\Delta S$, dihedrals | <u>$+ -0.313 \pm 0.004$</u> | <u>$+ -2.162 \pm 0.006$</u> | $\Delta\Delta G$ (Eqs. 3 & 2)<br>DOPC - DPPC |
| | $\Delta G$, sum, Eq. 3 | $7.83 \pm 0.08$ | $8.10 \pm 0.08$ | $-0.27 \pm 0.10$ |

Figure 4 illustrates some typical acyl chain conformations at the receptor surface. In some cases, a lipid has one acyl chain in contact with the protein surface and one in contact with other lipids. The chain that is in contact with the receptor shows increased gauche dihedrals, corresponding to a higher conformational entropy compared to acyl chains in the bulk, which are more ordered on average.

## Discussion

Previous analyses of all-atom simulations have characterized localized and specific lipid-GPCR interactions — sites on the protein surface which bind lipids at high affinity compared to the typical lipid-protein interaction (Song et al., 2019; Neale et al., 2015; Cang et al., 2013; Yen et al., 2018; Lee & Lyman, 2013; Lee et al., 2013; Rouviere et al., 2017; Yang & Lyman, 2019). This mode of interaction is simple to detect in simulations, since it yields long-lived lipid binding. It is straightforward to imagine how such interactions may have functional consequences (e.g., via allosteric coupling to ligand or G-protein binding), since they are similar to other small molecule-protein interactions.

Among MD studies which focus on cholesterol, Cang, et al. used density mapping around the *β*_2_-adrenergic receptor to show competitive and cooperative associations between 1-palmitoyl-2-oleoyl-glycero-3-phosphocholine (POPC) and cholesterol in 5-*μ*s all-atom simulations, and observed spontaneous binding of cholesterol at the cholesterol consensus motif (Cang et al., 2013). Lyman and coworkers (Rouviere et al., 2017; McGraw et al., 2019) reported a series of 6-*μ*s all-atom simulations of A_2A_R bound to either agonist or antagonist, also observing cholesterol binding to the cholesterol consensus motif, and reporting two new putative cholesterol binding sites. Several studies have characterized anionic lipid interactions with GPCRs at the inner leaflet, including close interactions with PIP_2_ that are elevated in the active, G-protein bound state and enhance receptor/G-protein interactions in some cases (Neale et al., 2015; Song et al., 2019; Yen et al., 2018). Relevant to the present work, Grossfield, Feller, and Pitman examined the role of direct interaction of polyunsaturated fatty acids (PUFA) with Rhodopsin, a class-A cousin of A_2A_R, using a series of short (100 ns) simulations to sample the lipid-protein interactions (Grossfield et al., 2006).

Conceptually contrasting with these previous reports, the present work addresses the functional role of preferential lipid solvation — lower affinity but still selective interactions between lipids and membrane proteins, that lead the protein to recruit a first shell composition that differs from the bulk. A recent paper by Yang and Lyman suggested enrichment of DOPC over DPPC at the surface of A_2A_R, but a lack of clear equilibration of the A_2A_R’s first lipid solvation shell hampered robust functional inferences (Yang & Lyman, 2019). These observations were based on long (15 μsec) all-atom simulations, which emphasized the need for an approach that obtains rigorous convergence of the lipid environment around the protein.

This work presents such an approach based on a coarse-grained transition matrix (CGTM) populated by a series of short simulations. The equilibrium distribution predicted by the CGTM yields the composition of the first solvation shell. Because the method is naively parallel, large scale computing resources can be deployed, and results obtained in a matter of days, without requiring specialized hardware or long, contiguous trajectories.

Analysis of the composition of the first solvation shell indicates that the receptor recruits DOPC over DPPC in both the inactive and partially-active states. More importantly, the free energy for DOPC to partition to the first solvation shell is significantly *more favorable* for the partially-active than for the inactive receptor, indicating a mechanism coupling receptor function and membrane composition that is elaborated below.

The state-dependent difference in receptor solvation suggests a functional mechanism based on lipid-dependent shifting of the R↔R* equilibrium, depicted schematically in Figure 5. The apo receptor fluctuates spontaneously between the inactive R state and the active R* state. (These spontaneous, thermally excited fluctuations underlie the observed constitutive activity of A_2A_R (McGraw et al., 2019), and many other receptors.) The fraction of receptors that are in R vs. R* is governed by the free energy difference between these two states:

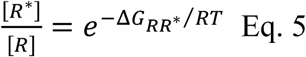

where 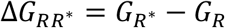 is the free energy difference between the R and R* states and [] denotes the fraction of receptors in the R or R* state. Since it is the R* state that catalyzes nucleotide exchange at the G-protein, this ratio determines the signaling output.

**Figure 5.**
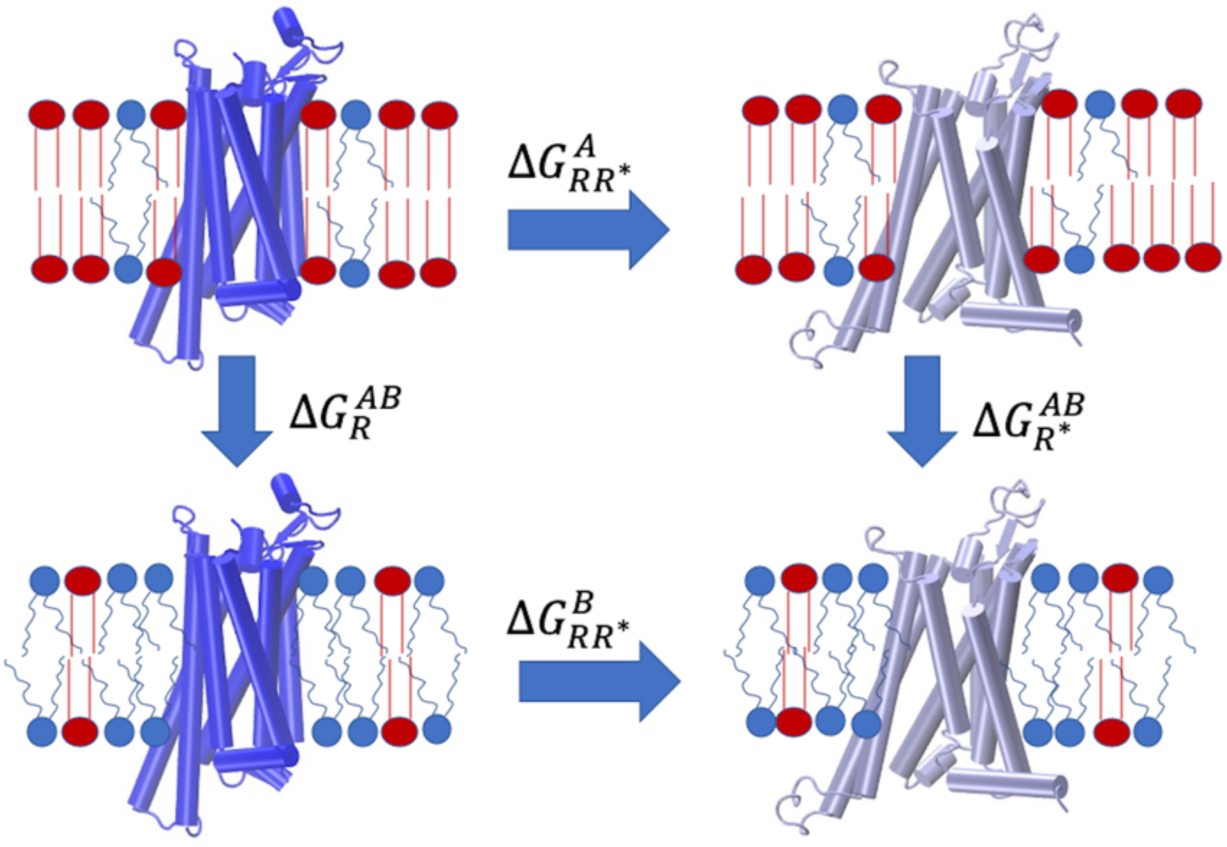
Activation energy depends on lipid solvation. In membrane composition “A,” the R→R* equilibrium (left to right arrow; inactive receptor in dark blue, active receptor in light blue) is determined by the free energy difference between R and R*, 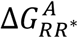, which differs in composition “B.” The vertical arrows indicate the free energy associated with a change in receptor environment in each state (e.g., as listed in Table 1). Differences in the R/R* equilibrium are observable as differences in constitutive activity and ligand affinities.

**Figure 6.**
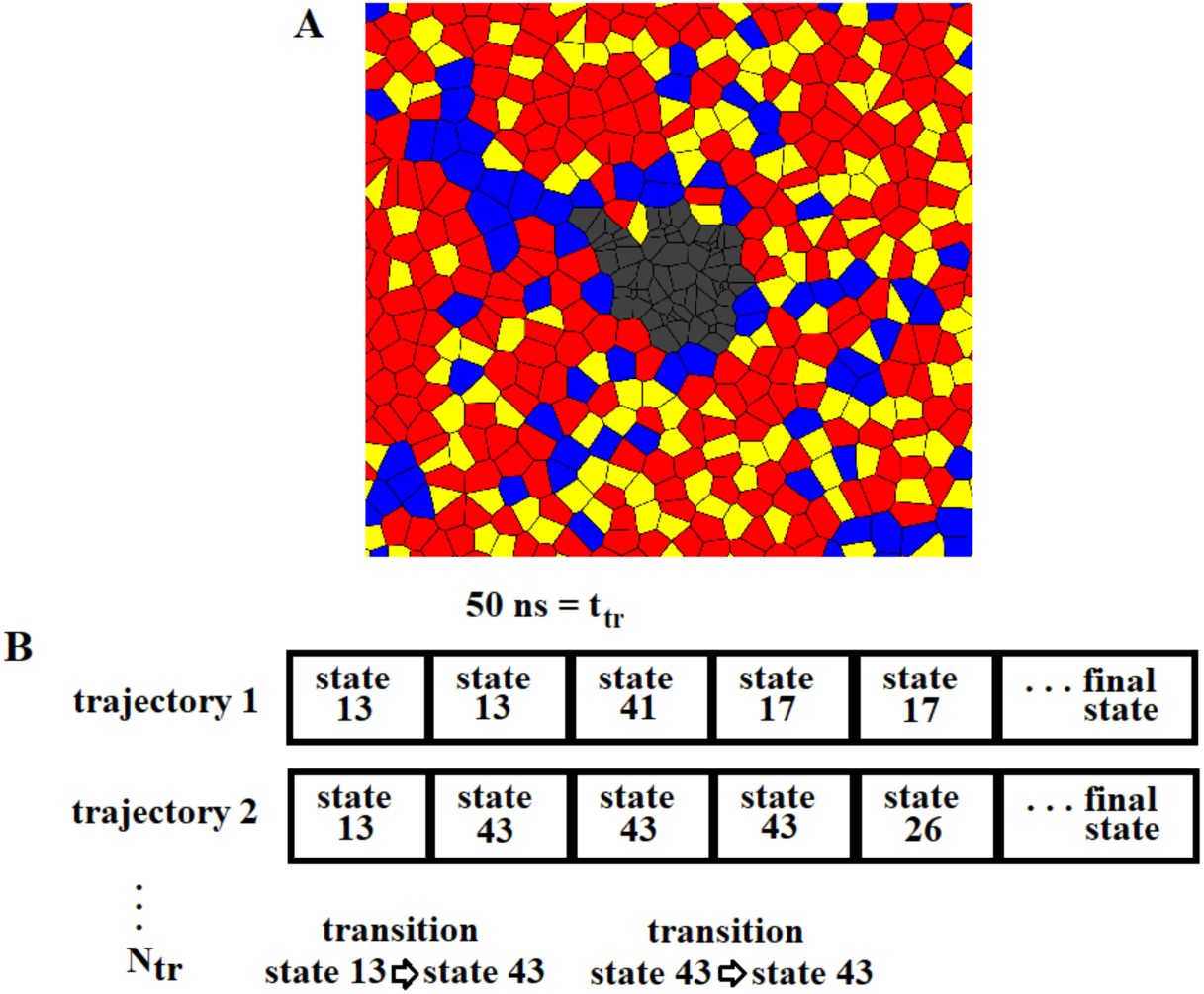
Method for constructing CGTM. Panel A: Example of Voronoi tessellation of the on lealeft. Voronoi cells for DOPC (blue), DPPC (red), and cholesterol (yellow). Agonist-bound receptor. Panel B: Schematic for definition of transitions used to populate the CGTM. Short simulations were initialized in states selected to span configuration space. Here, “state 13” is the initial state. Transitions were tallied with inclusion of redundancy. To test convergence, the number of replicas in each initial state (*N*_tr_) was varied, as well as the length of the trajectories (*t*_tr_).

The results presented here indicate that 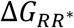 is lipid dependent, such that the R↔R* equilibrium is modulated by the chemical potential of lipids in the bulk. Consider a change in lipid composition which increases the chemical potential of unsaturated relative to saturated chains. This will favor receptor activation, as it drives unsaturated chains to the receptor surface. Because the R↔R* equilibrium directly impacts experimental observables, the proposed model is experimentally testable. Lipid mixtures that favor R* should increase constitutive signaling and apparent affinity of agonists, while mixtures that shift the equilibrium toward R will suppress constitutive activity and increase the affinity of inverse agonists.

Although the results presented herein are specific to the particular lipid mixture that was simulated, they establish a new conceptual framework for thinking about lipid-dependent membrane protein function. It is not possible to determine how specific effects on activity might change for a different lipid environment without performing simulations in that mixture — this is because the free energy to partition to the receptor surface depends on the chemical potential for each species in the bulk, and lipid chemical potentials are collective properties, determined by the overall composition. However, this challenge is readily overcome using the CGTM approach, which permits rapid sampling of the lipid environment around the receptor. Thus, the results can be easily extended to more realistic mixtures and to other integral membrane proteins. Comparison across systems, especially GPCRs (Sejdiu & Tieleman, 2020), should help to determine generic aspects of lipid-dependent GPCR function.

## Materials and Methods

### Determining the lipid solvation of A_2A_R

Lipids were assigned to a solvation shell around A_2A_R by a Voronoi tessellation based on centers of mass of the lipids and *α*-carbon atoms of the protein, following Beaven, et al (Beaven et al., 2017). Each lipid was assigned a shell number according to whether it shares a boundary with the receptor (first solvation shell), with a first-shell lipid (second solvation shell), and so on. Lipids in the fifth shell were considered “bulk” because this shell was furthest from the receptor and also contained ratios of lipid types nearest to the whole system averages (within 1%). Because site-specific binding was not considered, both leaflets were averaged together. Figure 6A shows an example of Voronoi boundaries for a single leaflet in a single frame.

The lipid composition of the first shell was characterized by the percent of each lipid type, rounded to the nearest 5% and constrained to total 100%. For example, a first-shell solvation state could be DPPC/DOPC/Cholesterol in a ratio of 0.35/0.45/0.20.

Because this binning does not consider the lipids’ spatial distribution, the resulting states are coarse grained. The procedure followed is identical to constructing a Markov State Model, but with coarse graining; hence the name Coarse-Grained Transition Matrix (CGTM) modeling. To compute the equilibrium distribution of states (*π*^Eq^), a CGTM was populated by a series of short (50 nsec) trajectories initialized in states selected to span the configuration space. For this analysis, 16 initial states were selected, and replicas were run in each of the 16 states for each receptor state (partially active and inactive). MD simulation details for these short simulations are described in Materials and Methods. Initial configurations were taken from the 15-*μ*sec simulations for under-sampled states. The more populated states were adequately represented in the original 15-*μ*sec simulations; therefore, nonoverlapping segments of the original trajectories were extracted.

Figure 6B illustrates how transitions between states were counted and the CGTM populated. Assuming the system can occupy a state *s*_*i*_ existing in *s* = [*s*_1_, *s*_2_ … *s*_N_], discrete sampling of the system results in a probability distribution *π* = [*π*_1_, *π*_2_ … *π*_N_] in which *π*_*i*_ is the sampled probability of the system occupying *s*_*i*_. Transition probabilities between states are computed from transition events *c*_*ij*_ between *s*_*i*_ and *s*_*j*_:

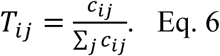

A CGTM 𝕋 containing these probabilities describes the evolution of state populations *π*_*i*_ over time:

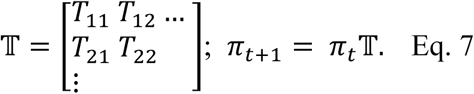

When there is no net population flow among states, the system is in equilibrium. *π*^Eq^ can be computed by solving the matrix equation:

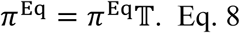

If there is no net probability flow among states, Eq. 8 holds and the system is in equilibrium. This is guaranteed to be true if microscopic detailed balance is satisfied:

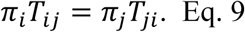

Detailed balance is therefore a sufficient condition for a system to exhibit equilibrium. Detailed balance implies “coarse balance”—balanced flux between every pair of arbitrarily defined set of microstates (Zuckerman, 2010). While coarse graining configuration space results in loss of the Markov property, the coarse balance property implies the reliability of equilibrium properties estimated using trajectories that sample the equilibrium probability distribution. Further verification of the computed *π*^Eq^ is therefore provided by checking the system for detailed balance and showing the robustness of *π*^Eq^ as a function of lagtime (time between discrete sampling events).

### Convergence of lipid distribution obtained by the CGTM approach

To test convergence of *π*^Eq^ computed from the CGTM, the number of replicas of short simulations in a given initial state (*N*_tr_) was varied from 5 – 9 and the trajectory length (*t*_tr_) was varied from 10 – 50 ns. The change in the cumulative root mean squared deviation of the state probabilities were computed:

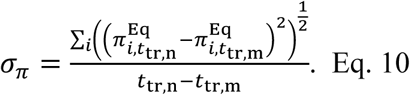

Here, 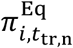 is the equilibrium probability of state *i* estimated from trajectory length *t*_tr,n_ in ns, with *t*_tr,n_ > *t*_tr,m_. Convergence of *σ*_*π*_ was compared for various values of *N*_tr_.

Supplemental Figure 6-1 shows that the state equilibrium probabilities are well converged using the CGTM approach. By the time each trajectory has reached 30 nsec the convergence metric *σ*_*π*_ shows that the cumulative state probabilities are changing by about 1%, and including more data (whether in the form of longer individual trajectories or as additional trajectories) does not improve the statistics. Supplemental Figure 6-1 shows that well-converged lipid solvation can be obtained for this system with a total of only 2.4 *μ*sec of total simulation time: (number of initial states)* *N*_tr_\**t*_tr_ = 16*5*30 ns = 2.4 *μ*s. This is significantly less than the duration of the initial 15 *μ*sec simulations which never reach clear convergence. Moreover, the CGTM approach is naively parallel — all data used to produce Figure 2 can easily be obtained in a single day on a commodity supercomputing resource.

While 5 trajectories/initial configuration shows good convergence of *π*^Eq^, 9 total replicas were run for each initial configuration. Supplemental Figure 6-2A shows the error in *π*^Eq^ computed among blocks of 3 trajectories each; actual error is likely considerably lower, since better convergence is obtained when all 9 trajectories are used to compute *π*^Eq^. Additionally, *T* and the resulting *π*^Eq^ computed for the agonist- and antagonist-bound receptors were checked for detailed balance. Deviation from Eq. 9 can be computed for each 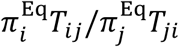 pair: aBs 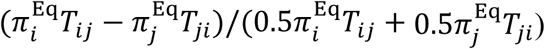. On average, observed transitions deviated from detailed balance by 5%. Detailed balance was robustly observed for high frequency transitions, with larger deviations seen for infrequent transitions.

Finally, the computed *π*^Eq^ is independent of lagtime, a necessary but not sufficient condition of a system in equilibrium (Swope et al., 2004). Intuitively, this can be understood as the system exhibiting predictable flow from one state to another despite passing through intermediate states for which the Markov property holds; meaning transitions into and out of those states are also well sampled. While *π*^Eq^ shown in Figure 2 are computed with a lagtime of 240 ps (the time between subsequent trajectory frames), increasing the lagtime to 960 ps shows no noticeable change in *π*^Eq^; the distributions overlap (Supplemental Figure 6-2B).

### Computation and convergence of configurational entropy of acyl chains

A free energy scan about each type of dihedral in the carbon backbone of DOPC and DPPC acyl chains is shown in Supplemental Figure 4-1. DOPC chains contain 3 types of rotatable dihedrals: C-C-C-C, C=C-C-C/C-C-C=C, and C-C=C-C. DPPC acyl chains have one type: C-C-C-C. Relative energy maxima from these plots were taken as boundaries for binning into dihedral states.

Supplemental Figure 4-2 compares the resulting acyl backbone dihedral *S* of 1^st^ shell and bulk lipids. For both lipid types, configurational entropy of acyl chains is greater at the protein surface than in bulk. Increase in *S* indicates it is entropically favorable for the acyl chains to neighbor the protein surface because they sample a larger dihedral configuration space. The acyl chains of saturated DPPC, which in the cholesterol-rich bulk are relatively ordered, exhibit the largest change in *S* moving from bulk to the protein surface. Therefore, while *S* contributes favorably to the solvation of the protein by both lipid types, it is entropically more favorable for saturated DPPC to solvate the protein.

Each DOPC acyl chain backbone contains 15 dihedrals, and each DPPC backbone 13. The dependence of *S* on the number of dihedrals considered to be coupled when defining a microstate is plotted in Supplemental Figure 4-3A. Accounting for correlations between dihedral states in the same lipid tail results in a decrease in the estimate of *S*, as expected — correlation reduces entropy. Notably, estimates of *S* for the DPPC backbone do not decrease when coupling beyond 1 or 2 neighboring dihedrals (N = 2 or N = 3) is considered. However, because the DOPC acyl backbone contains multiple types of dihedrals which are each assigned to distinct states and can occupy only specific sites, it is necessary to consider the entire tail to be coupled to get an accurate estimate of *S* for DOPC.

Supplemental Figure 4-3B shows the convergence of *S* with increasing sample size. Each point represents the average *S* computed from different sized non-overlapping trajectory blocks. Good convergence is seen for both lipid types. Convergence is reached more rapidly if fewer neighboring dihedrals are considered to be coupled because this reduces the number of possible states. An upper limit of *S* can therefore be computed, but given the good convergence seen in Figure 4-3B, the present results consider the entire tail to be coupled.

### Molecular dynamics simulations

Initial coordinates for the inactive state of the receptor were based on the 1.8 Å resolution structure bound to ZM241385, Protein Data Bank (PDB) entry <u>4EIY</u> (Liu et al., 2012). The partially active state was based on the UK432097-bound structure in which intracellular loop 3 is replaced with a BRIL fusion construct, PDB entry <u>3QAK</u>(Xu et al., 2011); the BRIL construct was excised, and the missing residues were rebuilt as described previously (Lee et al., 2013). The protein was embedded in a symmetric lipid bilayer containing 504 DPPC, 273 cholesterol, and 132 DOPC molecules and solvated with approximately 55000 modified TIP3P waters (Jorgensen et al., 1983; Durell et al., 1994), one Na^+^ as observed in the crystal structure, and 10 Cl^-^ to neutralize the system. The lipid mixture (0.55:0.15:0.30 DPPC:DOPC:Chol) is in the liquid–liquid coexistence region of the phase diagram. The protein and lipids were modeled with the CHARMM36 force field (Klauda et al., 2010; Huang et al., 2017; Watts et al., 2018), and the ligand was modeled with the CHARMM general force field (Vanommeslaeghe et al., 2010) with atoms typed by the ParamChem server (Vanommeslaeghe & MacKerell, 2012).

Production simulations were run on Anton2. Integration was performed under constant pressure (1 atm), temperature (295 K), and particle number with the multigrator (Lippert et al., 2013) method with a 2.5 fsec time step. Temperature was controlled by a Nose-Hoover (Nosé, 1984) thermostat coupled every 24 time steps and pressure controlled by the Martyna–Tobias–Klein barostat (semi-isotropic) coupled every 480 time steps (Martyna et al., 1994). Electrostatics were computed using the *k*-space Gaussian split Ewald method (Shan et al., 2005) with long-range interactions computed every third time step and hydrogens constrained by M-SHAKE (Kräutler et al., 2001). Lennard-Jones interactions were cut off at ≥11 Å. Production simulations were run for 15 *μ*sec, and configurations were stored every 240 psec.

Additionally, to verify the equilibrium lipid solvation of A_2A_R, a series of 50-nsec simulations (“short simulations”) were run using GROMACS 2019.4 simulation package (Abraham et al., 2015). The simulations were performed in the NPT ensemble at 1 bar and 295 K with the Nosé−Hoover thermostat (Hoover, 1985; Nosé, 1984; Nosé, 1984) and Parrinello−Rahman barostat (Parrinello & Rahman, 1980; Parrinello & Rahman, 1981). A coupling constant of τ_*p*_ = 5.0 psec was used for pressure control, and for temperature τ_*t*_ = 1 psec. The protein, lipid bilayer, and water were separately coupled to the heat bath, and the semi-isotropic pressure coupling was applied separately in the *xy*-direction (bilayer plane) and the *z*-direction (bilayer normal). Electrostatics were computed with the particle-mesh Ewald method with a real-space cutoff of 9 Å, Lennard-Jones interactions were truncated at 11 Å, and configurations were stored every 240 psec.

Images of lipids and receptors were generated and values of acyl chain dihedrals determined with the Visual Molecular Dynamics program (Humphrey et al., 1996). Internal energies and nonbonded interaction energies were computed using the CHARMM program (Brooks et al., 2009).

## Acknowledgements

The authors thank Alex Sodt and Dan Zuckerman for extensive discussion regarding implementation of the CGTM, and Alex for assistance with Voronoi analysis. This work used the Extreme Science and Engineering Discovery Environment (XSEDE) supported by National Science Foundation Grant ACI-1548562. Anton2 computer time was provided by the Pittsburgh Supercomputing Center (PSC) through Grant R01GM116961 from the National Institutes of Health. The Anton2 machine was made available by D. E. Shaw Research

---

**Supplemental Figure 6-1.**
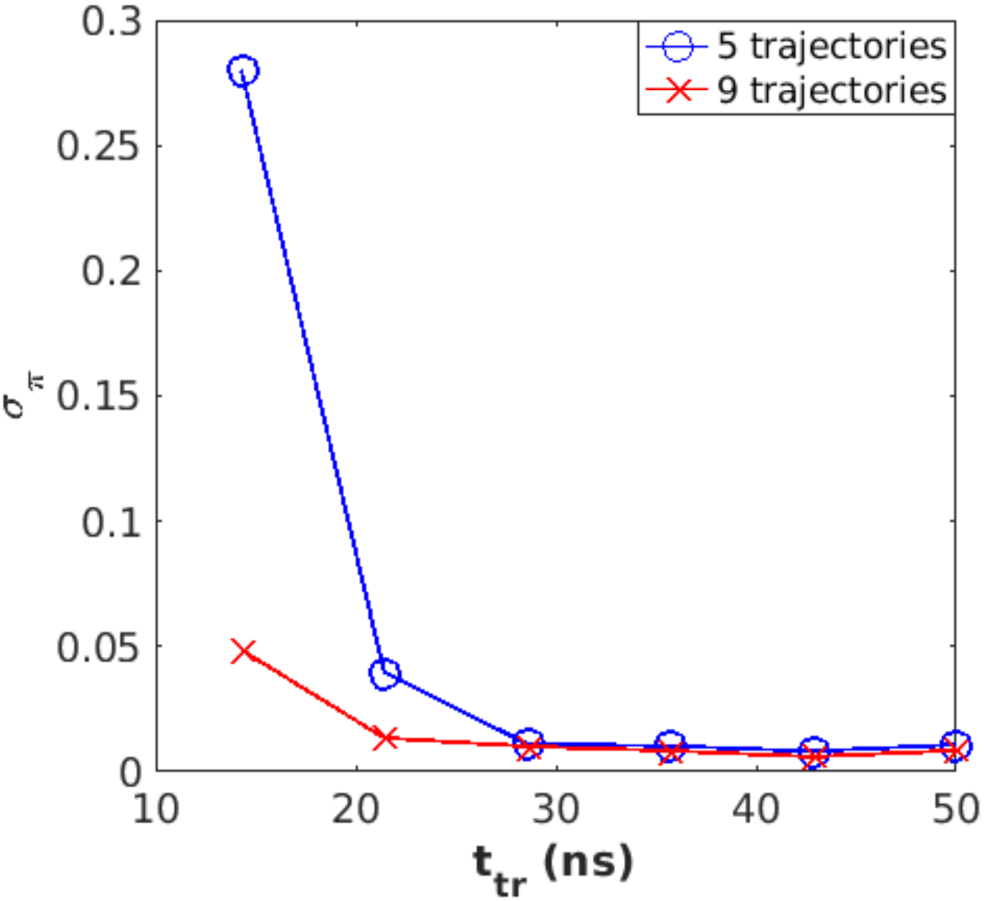
Convergence of *π*^Eq^ is achieved with modest sampling. The convergence metric (*σ*_*π*_, Eq. 10) demonstrates that the state probabilities are well converged after each individual trajectory reaches 30 nsec, using 5 independent trajectories per initial state. Blue: 5 trajectories per initial state. Red: 9 trajectories per initial state. Agonist-bound receptor.

**Supplemental Figure 6-2.**
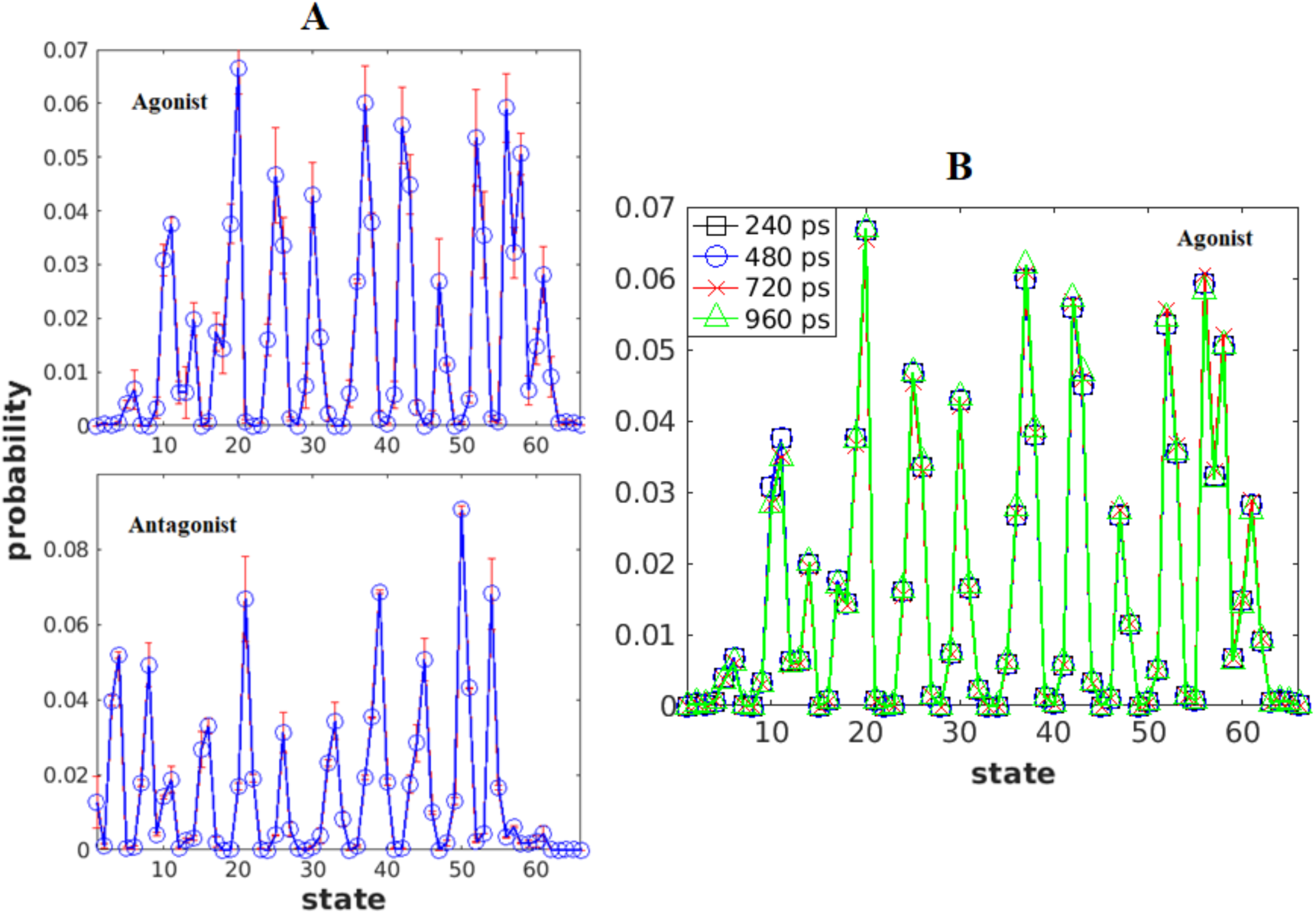
Statistical reliability of *π*^Eq^. Panel A: Error in *π*^Eq^ computed by block averaging. Standard error in *π*^Eq^ computed from 3 sets of 3 trajectories each per initial configuration for the partially active state (top) and inactive state (bottom). Panel B: Equivalence of *π*^Eq^ computed using different lagtimes. Only slight variation in state probabilities are seen if *π*^Eq^ is computed with a lagtime of 240, 480, 720, or 960 ps. *π*^Eq^ is independent of lagtime in equilibrated systems. Agonist receptor; computed with overlapping trajectory windows. Similar equivalence was exhibited by antagonist-bound receptor.

**Supplemental Figure 4-1.**
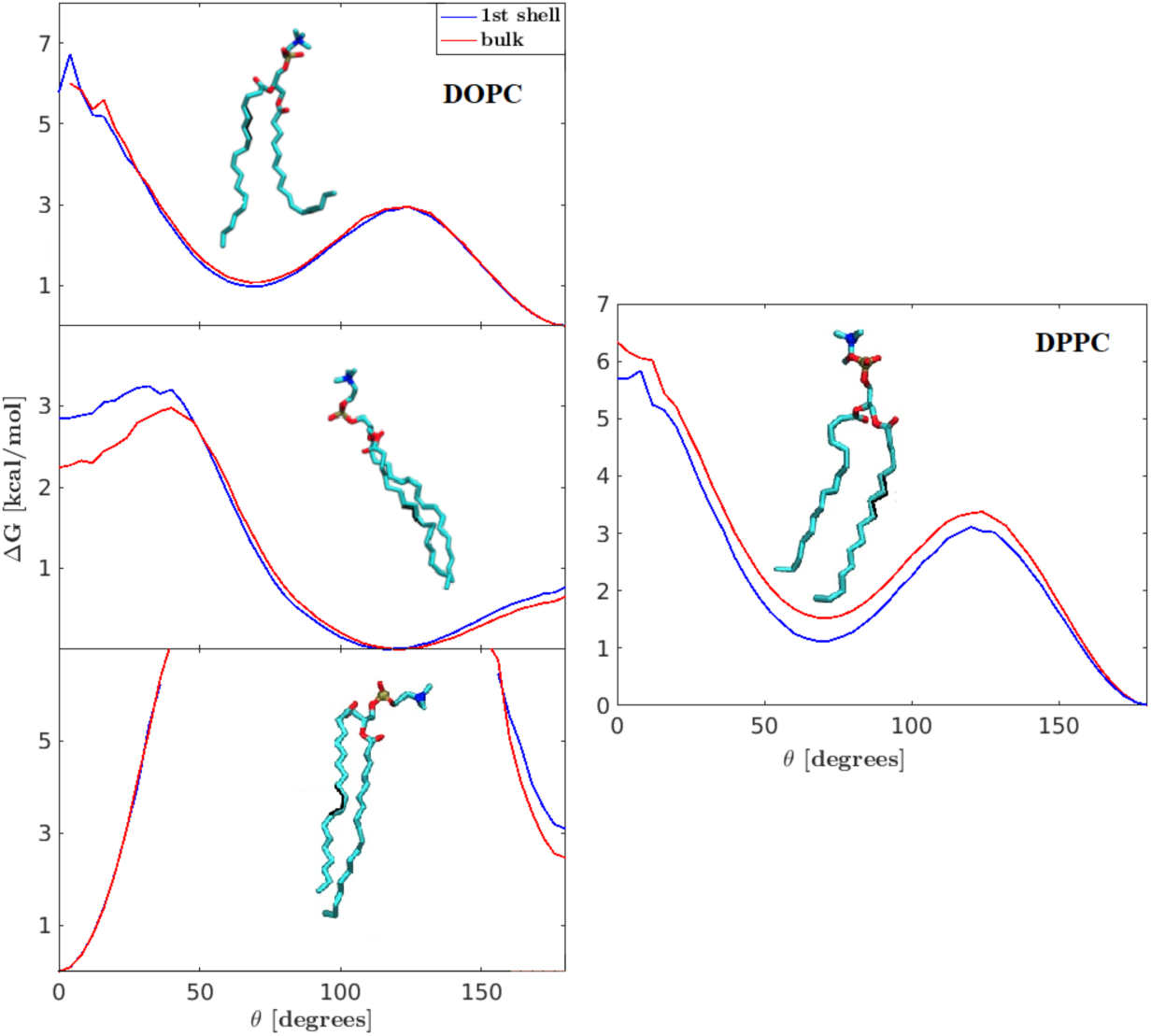
DOPC and DPPC free energy landscapes for various dihedral types. Dihedral types in acyl chains; example minimum energy configurations shown with dihedrals highlighted in black. (left) DOPC dihedrals include (top) C-C-C-C, maxima at 0 and ±130°; (middle) C-C-C=C, maxima at 180 and ±45°; and (bottom) C-C=C-C, maxima at ±100°. (right) DPPC includes C-C-C-C, maxima at 0 and ±130°.

**Supplemental Figure 4-2.**
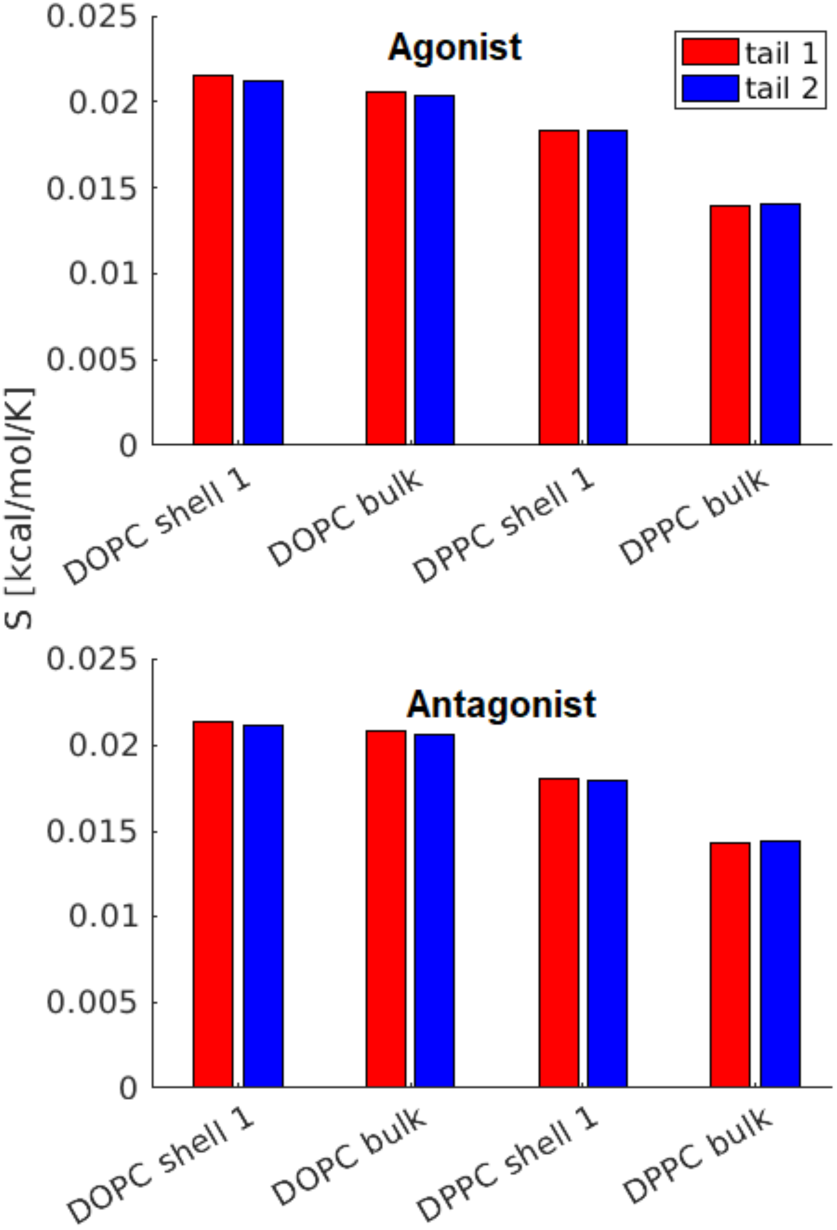
Acyl chain dihedral entropy (*S*) of first shell and bulk lipids. Error computed from block averaging is on the order of 1%; therefore, error bars are not shown. For both receptors, entropy favors DPPC partitioning from bulk to the first shell significantly more than DOPC.

**Supplemental Figure 4-3.**
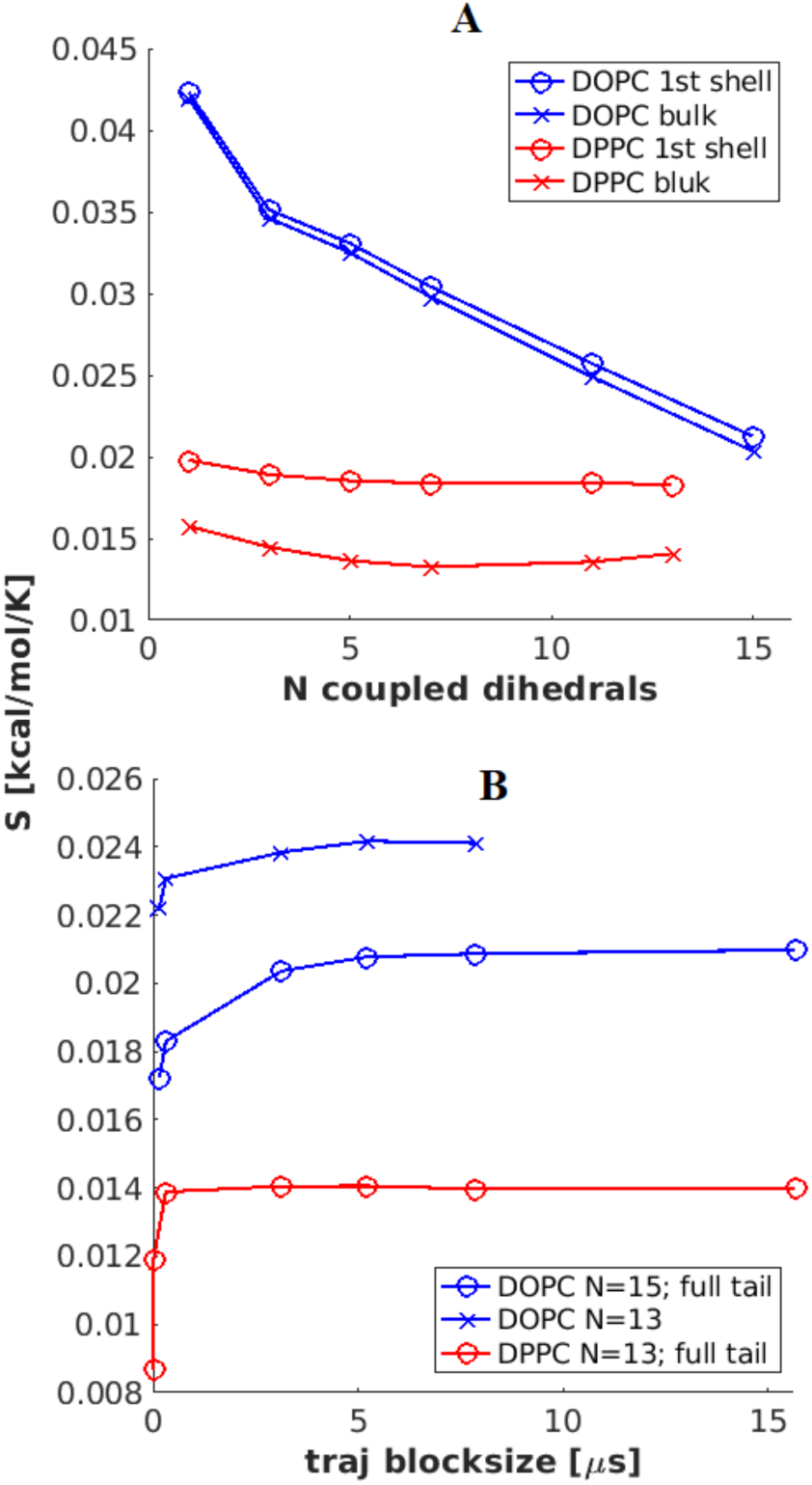
Convergence of acyl backbone dihedral entropy (*S*). Panel A: Dependence of *S* on the number of neighboring dihedrals (*N*) considered to be coupled. Sampled from simulation of agonist-bound receptor, *sn*-2 chain. Error computed from block averaging is on the order of 1%; therefore error bars are not shown. Panel B: Convergence of *S* as a function of trajectory length. Plotted as a function of length of trajectory used in block averaging. Two data sets are included for DOPC, one with the entire tail considered to be coupled, and one with 13 of the 15 dihedrals considered to be coupled; *N* = 13 or 15 respectively. Sampled from simulation of agonist-bound receptor, *sn*-1 chain. Error computed from block averaging is on the order of 1%; therefore error bars are not shown.

